# Differential regional decline in dopamine receptor availability across adulthood: Linear and nonlinear effects of age

**DOI:** 10.1101/358200

**Authors:** Kendra L. Seaman, Christopher T. Smith, Eric J. Juarez, Linh C. Dang, Jaime J. Castrellon, Leah L. Burgess, M. Danica San Juan, Paul M. Kundzicz, Ronald L. Cowan, David H. Zald, Gregory R. Samanez-Larkin

## Abstract

Theories of adult brain development, based on neuropsychological test results and structural neuroimaging, suggest differential rates of age-related change in function across cortical and subcortical sub-regions. However, it remains unclear if these trends also extend to the aging dopamine system. Here we examined cross-sectional adult age differences in estimates of D2-like receptor binding potential across several cortical and subcortical brain regions using PET imaging and the radiotracer [18F]fallypride in two samples of healthy human adults (combined *N* = 132). After accounting for regional differences in overall radioligand binding, estimated percent difference in receptor binding potential by decade (linear effects) were highest in most temporal and frontal cortical regions (∼6–16% per decade), moderate in parahippocampal gyrus, pregenual frontal cortex, fusiform gyrus, caudate, putamen, thalamus, and amygdala (∼3–5%), and weakest in subcallosal frontal cortex, ventral striatum, pallidum, and hippocampus (∼0–2%). Some regions showed linear effects of age while many showed curvilinear effects such that binding potential declined from young adulthood to middle age and then was relatively stable until old age. Overall, these data indicate that the rate and pattern of decline in D2 receptor availability is regionally heterogeneous. However, the differences across regions were challenging to organize within existing theories of brain development and did not show the same pattern of regional change that has been observed in gray matter volume, white matter integrity, or cognitive performance. This variation suggests that existing theories of adult brain development may need to be modified to better account for the spatial dynamics of dopaminergic system aging.

## Introduction

Studies of neurocognitive function, as well as gray matter and white matter structure, have identified differential decline in aging. Some theories of adult brain development suggest that compared to other brain regions, the frontal lobes show steeper age-related decline. These theories include the frontal lobe hypothesis of cognitive aging (West, 1996, 2000) and speculation about an anterior-posterior gradient based on studies of both gray matter volume (e.g. (Raz & Rodrigue, 2006)) and white matter integrity (e.g. (Sullivan & Pfefferbaum, 2006)). Recent work, however, has suggested a more nuanced view of adult development and aging. For instance, cognitive functions that are thought to be independent of the frontal lobes also display significant age-related decline (Greenwood, 2000; Rubin, 1999) while tasks that are assumed to be frontal-dependent can be impaired with caudate or putamen lesions (Rubin, 1999). Further, several studies have shown that a monotonic decrease in white matter integrity extends well beyond the frontal lobes (Bennett, Madden, Vaidya, Howard, & Howard, 2010; Davis et al., 2009).

These qualifiers led to revised theories of adult brain development, which suggest that relative to medial brain regions, lateral regions undergo greater age-related decline. For example, the dorsolateral prefrontal theory of cognitive aging suggests that compared to ventromedial-dependent tasks, greater age-related differences are found in dorsolateral-dependent tasks (MacPherson, Phillips, & Della Sala, 2002). Similarly, the “last-in, first-out” or retrogenesis hypothesis of aging (Davis et al., 2009; Fjell et al., 2009; Raz, 2000) suggests regions which mature later in development and evolution are the first to display age-related vulnerability to decline, while phylogenetically older areas of the brain are preserved. This theory is supported by studies showing age-related gray matter decreases in the association cortices in middle age (McGinnis, Brickhouse, Pascual, & Dickerson, 2011), medial temporal lobes in early-old age (Fjell et al., 2013; Raz, Ghisletta, Rodrigue, Kennedy, & Lindenberger, 2010; Raz et al., 2005; Yang et al., 2016), and primary sensory cortices later in late-old age (Yang et al., 2016). Additionally, some studies have shown that in healthy aging, the hippocampus is best fit by a quadratic model because it is relatively preserved until fairly late in the adult life span (Fjell et al., 2014; Fjell et al., 2013). Thus, theories of adult brain development, supported by neuropsychological and structural neuroimaging evidence, suggest differential rates of age-related change across cortical and subcortical sub-regions.

Given these differential rates of age-related change in brain structure and cognitive function, it is possible that there are also differential rates of age-related change in dopaminergic function across cortical and subcortical regions. The vast majority of studies examining the correlations between age and dopamine have reported linear declines in non-displaceable binding potential (BP_ND_) in D2-like receptors in striatal regions (e.g. (Bäckman & Farde, 2001; Inoue et al., 2001)), with only a handful of studies examining BP_ND_ in frontal regions (e.g. (Kaasinen et al., 2000; Ouchi et al., 1999)). Specifically, studies of D2-like receptor BP_ND_ report wide-ranging age-related effects, ranging from slightly negative (Kim et al., 2011) to strongly negative (Bäckman et al., 2000; Mukherjee et al., 2002). A recent meta-analysis showed strongly negative linear effects of adult age on D2-like receptors in both frontal (*r* = –.66) and striatal (*r* =–.54) regions (Karrer, Josef, Mata, Morris, & Samanez-Larkin, 2017). In exploratory analyses, this meta-analysis also showed that linear and quadratic effects of age fit the data equally well. However, it is difficult to rule out potential contributions of methodological or sample differences across studies to these meta-analytic observations. Only a few individual studies have examined nonlinear effects of age on dopamine measures and these studies reported concave-down quadratic effects of age on dopamine transporters (Mozley et al., 1996; van Dyck et al., 2002). However, non-linear effects have not yet been systematically investigated in individual studies of D2-like receptors.

In part because of the limited ability for D2 radiotracers (with varying affinities) to capture receptor availability in both receptor dense (striatum) and sparse (frontal cortex) areas in the same scan session and the relatively low spatial resolution of many PET scanners, the vast majority of prior studies of age differences in the dopamine system have used large regions of interest (ROIs) spanning the whole frontal lobe or striatum. Few studies have examined the potential differences between sub-regions within these relatively large structures. Further, these prior studies did not use partial volume correction, which given the differential rates of age-related gray matter atrophy, could impact BP_ND_ levels (Smith et al., 2017).

Thus, in the present study we examined age effects in regional dopamine BP_ND_ across adulthood in cortical and subcortical sub-regions. We examined partial volume corrected (PVC) dopamine BP_ND_ of D2-like receptors in two cross-sectional, adult life-span studies. Using [^18^F]Fallypride, which provides broad coverage throughout both cortical and subcortical regions (Slifstein et al., 2004), allowed us to explore regional age differences in the dopamine system across the brain. Study 1 (*N* = 84) included participants continuously sampled across the adult life span and Study 2 (*N* = 48) included a group design with younger adults and older middle-aged adults. On the basis of theories of adult brain development described above, we hypothesized that BP_ND_ in most regions would show strong, negative effects of age, with steeper declines in lateral and frontal regions than in medial and posterior regions. There is also some limited evidence for linear declines in dopaminergic function with age in the hippocampus (Kaasinen et al., 2000; Stemmelin, Lazarus, Cassel, Kelche, & Cassel, 2000). However, based on anatomical studies, we hypothesized that BP_ND_ in the hippocampus would display preservation across most of adulthood with accelerated decline in old age.

## Methods

Both data sets (Study 1 and Study 2) were collected as part of large-scale multimodal neuroimaging projects focused on decision making. Subsets of the Study 1 behavioral (Seaman et al., 2016), fMRI (Seaman et al., 2018) and PET (Dang et al., 2017; Dang et al., 2016; Smith et al., 2017) data were previously included in other publications. Specifically, age effects on D2- like BP_ND_ in a subset of Study 1 participants were reported or noted in three previous publications (Dang et al., 2017; Dang et al., 2016; Smith et al., 2017). However, these were limited to non-PVC striatal ROIs (Dang et al., 2017; Dang et al., 2016) or very large cortical ROIs (i.e., frontal cortex, parietal cortex) that averaged across all gyri within a lobe (Smith et al., 2017). Here we focus on *regional* age differences in partial-volume corrected D2-like receptor BP_ND_ across the adult life span using the full sample from Study 1 (not previously reported) and a new study (Study 2).

### Participants

For both studies, volunteers were recruited from the Nashville community for a multiday, multimodal neuroimaging study of decision making using the Vanderbilt School of Medicine subject database of healthy adults, Research Match (www.researchmatch.org), and a combination of newspaper, radio, and local TV advertisements. All participants were mentally and physically healthy; exclusion criteria included a history of psychiatric illness, head trauma, any significant medical condition, pregnancy, substance abuse, or any condition that would interfere with MRI (e.g. claustrophobia or metal implants). For Study 1, of the 92 adult volunteers recruited, a total of 84 participants (*M* = 49.43, Range = 22 to 83 years old) completed both MRI and PET scans. For Study 2, of the 73 volunteers recruited, 48 participants (*M* = 41.40, Range = 20−65 years old) completed MRI and baseline PET scans. Study 2 participants additionally completed a second [18F]fallypride PET scan after taking oral d-amphetamine to measure dopamine release, and a third PET scan using [18F]FE-PE2I to measure dopamine transporter availability. Thus, the high rate of exclusions in Study 2 in part reflects stringent criteria regarding blood pressure and the time commitment necessary to complete the study. The dopamine release and transporter data are not included here. All participants gave written informed consent and were compensated $350 for Study 1 and $370-675 depending on (1) task performance, (2) the number of PET scans completed and (3) time spent on the study for Study 2. Approval for all methods was obtained from the Vanderbilt University Human Research Protection Program and the Radioactive Drug Research Committee.

### Cognitive Assessment

Participants completed a battery of neuropsychological assessments during a separate session. Mean performance on this test battery, as well as the correlation of each measure with age and/or the difference between age groups, are displayed in Table 1. All participants in both studies displayed normal performance on cognitive tests. In Study 1 we found the expected age effects in measures of fluid intelligence (e.g. Digit Span, Numeracy, and Delayed Recall) and maintenance of crystallized intelligence (e.g. Vocabulary) across the adult life span. In Study 2, there was only a significant difference in delayed recall between younger and middle-aged adults; there were no other group differences in cognitive performance.

**Table 1.** Participant *Characteristics*

| Variable | Study 1 (N=84) |  | Study 2 (N=48) |  |  |
| --- | --- | --- | --- | --- | --- |
|  | M (SD) | *r* [95% CI] with age | YA<br>M (SD) | MA<br>M (SD) | Group t-value<br>[95% CI] |
| Age | 49.43 (17.64) |  | 25.78 (2.61) | 55.65 (3.72) |  |
| Gender | 48F/36M |  | 12F/11M | 10F/10M |  |
| Digit Span<sup>a</sup> | 16.12 (3.96) | **-0.28 [-0.47, -0.07]** | 20.52 (3.95) | 21.85 (4.13) | 1.07 [-1.18 , 3.83] |
| Numeracy<sup>a</sup> | 11.81 (3.24) | **-0.27 [-0.46, -0.06]** | 12.52 (1.5) | 15.35 (7.88) | 1.58 [-0.9 , 6.56] |
| Paired Associates Delayed Recall<sup>b</sup> | 5.85 (2.34) | **-0.61 [-0.73, -0.45]** | 7.22 (1.62) | 5.95 (2.14) | **-2.16 [-2.46 , -0.08]** |
| Shipley Vocabulary Subscale<sup>a</sup> | 33.67 (5.39) | 0.15 [-0.07, 0.35] | 33.3 (3.46) | 32.1 (9) | -0.56 [-5.62 , 3.21] |

*Notes.* Digit Span and Paired Associates Delayed Recall from the WMS-III, Wechsler Memory Scale-Third Edition, (Wechsler, 1997); Numeracy, (Peters, Dieckmann, Dixon, Hibbard, & Mertz, 2007); Shipley Vocabulary Subscale, (Shipley, 1940); Trails Test, (Corrigan & Hinkeldey, 1987).
YA = Younger adults; MA = older Middle Aged adults
Significant correlations denoted in bold.
No significant group differences in Study 2.
<sup>a</sup>Digit Span, Vocabulary, and Numeracy were not recorded for one participant in Study 1.
<sup>b</sup>Delayed Recall not recorded for five participants in Study 1.

### PET data acquisition and processing

PET imaging was collected at Vanderbilt University Medical Center. [^18^F]Fallypride was produced by the PET radiochemistry laboratory following the synthesis and quality control guidelines described in US Food and Drug Administration IND 47,245. A 5.0 mCi slow bolus injection of [18F]Fallypride was followed by three, 3D emission scans in a GE Discovery STE scanner (3.25 mm axial slices with in-plane pixel dimensions of 2.3 × 2.3 mm). The same scanner was used for both studies. Prior to each emission scan, CT scans were collected for attenuation correction. Scanning lasted for approximately 3.5 hours, with two 15-minute breaks for participant comfort. Decay, attenuation, motion, and partial volume correction was performed on the PET scans and voxelwise BP_ND_ maps, which represent the ratio of specifically-bound [^18^F]Fallypride to its free concentration, were calculated using the PMOD Biomedical Imaging Quantification software (see (Dang et al., 2016; Smith et al., 2017) for greater detail).

### MRI data acquisition

Structural MRI scans were collected using a 3-T Phillips Intera Achieva MRI scanner using a 32-channel head coil. T1- weighted high-resolution anatomical scans (repetition time = 8.9 ms, echo time = 4.6 ms, field of view = 256 × 256, voxel dimensions = 1 × 1×1 mm) were obtained for each participant. These structural scans facilitated co-registration and spatial normalization of the PET data.

### Partial volume correction

Using the Hammers atlas (Gousias et al., 2008; Hammers et al., 2003), both MRI and PET data were parcellated into 62 bilateral cortical, 12 bilateral subcortical, 3 posterior fossa, 5 ventricle, and 1 white matter regions of interest (a total of 83 regions). Following parcellation, the MRI and PET data were co-registered, PET data was resampled to MRI space, and then the partial volume correction (PVC) procedure available in PMOD’s PNEURO module was applied to the PET data. PNEURO uses the GTM method (Rousset, Collins, Rahmim, & Wong, 2008; Rousset, Ma, & Evans, 1998), which restricts PVC to the PET signal of structurally defined regions of interest. To evaluate the co-registration between the MRI and PET data, we calculated quality control metrics using the PFUS module in PMOD 3.9 for a subset of 42 participants, including the oldest 10 participants in each study. It is our experience that due to gray matter loss with age, the oldest subjects are usually the hardest to co-register. The average Dice coefficient between PET data warped to MRI space and the MRI data itself, which is a ratio of the number of true positives compared to the number of true positives plus the number of false positives, was 0.86 ± 0.02. This suggests that the registration methods were successful and consistent across the sample. Further, we tested whether there were any age differences in registration quality control metrics provided by PFUS (sensitivity, specificity, Jaccard index) across the subsamples and found no relationship between age and any measure of registration quality. Time activity curves (TACs) from each region were extracted from the PET data after PVC and fit with a simplified reference tissue model (Lammertsma & Hume, 1996) using PMOD’s PKIN module where a gray matter bilateral cerebellum ROI was used as the reference region (see Smith et al., 2017 for greater detail).

### Regions of Interest

Prior to analysis, brainstem, white matter, occipital lobes, and ventricles were excluded from consideration for analysis because these regions have no or very low levels of dopamine receptors. The cerebellum was also excluded because it was used as the reference region in the TAC modeling. Although BP_ND_ is low across many cortical regions, studies have documented that meaningful signal can be extracted from regions with low binding (Mukherjee et al., 2002). To further limit the number of analyses, for each region of interest (ROI) we calculated the bilateral average BP_ND_ within each participant, giving a total of 33 bilateral ROIs.. Within each study, we screened these bilateral BP_ND_ averages for outliers, cutting any values that were more than 1.5 times outside the interquartile range (Study 1: *M* = 3.15, Range = 0 to 17 participants excluded in each ROI, Study 2: *M* = 2.08, Range = 0 to 8 participants excluded in each ROI). Based on this screening, all 33 bilateral ROIs were retained for primary analyses. Pictures, scatterplots, and statistics for each ROI are available in an interactive app online at http://13.58.222.229:3838/agebp/. Data and code are available online at https://github.com/klsea/agebp or https://osf.io/h67k4/.

### Statistical Analyses

Despite the fact that the two studies were carried out by the same lab, using the same PET camera, scanning protocol, and preprocessing and analysis pipelines, the two studies differed significantly in average BP_ND_ in many regions (Table 2). This may reflect the fact that one study involved a placebo condition (Study 2, whereas the other did not), and there were subtle differences in recruitment mechanisms reflecting challenges recruiting a healthy middle aged population to participate in a multiday study. Because of this difference, our baseline model included study, along with sex, which has been suggested to affect D2 receptor availability (Pohjalainen, Rinne, Någren, SyvÄlahti, & Hietala, 1998), as control variables to ensure that these variables did not exert an influence on estimates of D2 declines with aging. Age effects were tested with linear and quadratic regressions carried out using the lm command in the R programming language.

**Table 2.** Mean (SD) for D2-like receptor availability (BP_ND_) across two studies using [18F]Fallypride PET. Significant differences are denoted in bold.

| Region | Overall<br>BPND M<br>(SD) | Study 1<br>BPND M<br>(SD) | Study 2<br>BPND M<br>(SD) | Study mean<br>difference [95%<br>CI] |
| --- | --- | --- | --- | --- |
| Ventral striatum | 37.16 ( 8.01 ) | 39.21 (7.66) | 33.33 (7.27) | **-5.89 [-8.6, -3.17]** |
| Putamen | 33.02 ( 4.87 ) | 32.7 (5.09) | 33.63 (4.41) | 0.93 [-0.77, 2.64] |
| Caudate nucleus | 26.9 ( 5.2 ) | 25.92 (5.15) | 28.64 (4.87) | **2.71 [0.91, 4.51]** |
| Pallidum | 14.83 ( 3.47 ) | 15.84 (3.28) | 13.13 (3.12) | **-2.72 [-3.86, -1.57]** |
| Subcallosal area | 9.23 ( 3.88 ) | 9.86 (3.79) | 8.03 (3.82) | **-1.83 [-3.27, -0.39]** |
| Amygdala | 3.02 ( 0.65 ) | 3.2 (0.64) | 2.69 (0.56) | **-0.51 [-0.72, -0.3]** |
| Insula | 2.44 ( 0.64 ) | 2.33 (0.57) | 2.64 (0.7) | **0.32 [0.08, 0.56]** |
| Thalamus | 2.43 ( 0.39 ) | 2.4 (0.34) | 2.47 (0.47) | 0.07 [-0.09, 0.22] |
| Anterior temporal lobe, lateral | 1.67 ( 0.48 ) | 1.56 (0.37) | 1.86 (0.59) | **0.3 [0.12, 0.49]** |
| Anterior temporal lobe, medial | 1.64 ( 0.45 ) | 1.5 (0.35) | 1.89 (0.5) | **0.39 [0.23, 0.56]** |
| Fusiform gyrus | 1.62 ( 0.39 ) | 1.55 (0.35) | 1.75 (0.44) | **0.19 [0.04, 0.35]** |
| Superior temporal gyrus, anterior | 1.44 ( 0.52 ) | 1.26 (0.4) | 1.75 (0.56) | **0.49 [0.3, 0.67]** |
| Middle and inferior temporal gyrus | 1.4 ( 0.45 ) | 1.29 (0.36) | 1.59 (0.54) | **0.31 [0.13, 0.48]** |
| Hippocampus | 1.34 ( 0.43 ) | 1.47 (0.44) | 1.11 (0.3) | **-0.36 [-0.48, -0.23]** |
| Straight gyrus | 1.23 ( 0.44 ) | 1.05 (0.34) | 1.52 (0.43) | **0.47 [0.32, 0.62]** |
| Medial orbital gyrus | 1.19 ( 0.38 ) | 1.03 (0.29) | 1.45 (0.38) | **0.42 [0.3, 0.55]** |
| Subgenual frontal cortex | 1.01 ( 0.55 ) | 0.72 (0.35) | 1.53 (0.43) | **0.81 [0.66, 0.97]** |
| Posterior orbital gyrus | 0.94 ( 0.39 ) | 0.78 (0.31) | 1.19 (0.37) | **0.41 [0.28, 0.54]** |
| Superior temporal gyrus, posterior | 0.82 ( 0.39 ) | 0.7 (0.32) | 1.05 (0.42) | **0.35 [0.21, 0.49]** |
| Posterior temporal lobe | 0.81 ( 0.35 ) | 0.72 (0.31) | 0.98 (0.35) | **0.25 [0.13, 0.38]** |
| Parahippocampal and ambient gyri | 0.8 ( 0.27 ) | 0.71 (0.18) | 0.96 (0.33) | **0.26 [0.15, 0.36]** |
| Lateral orbital gyrus | 0.64 ( 0.4 ) | 0.48 (0.25) | 0.93 (0.45) | **0.45 [0.3, 0.59]** |
| Anterior orbital gyrus | 0.63 ( 0.37 ) | 0.46 (0.25) | 0.93 (0.37) | **0.47 [0.35, 0.59]** |
| Inferiolateral remainder of parietal lobe | 0.61 ( 0.42 ) | 0.43 (0.3) | 0.92 (0.42) | **0.49 [0.35, 0.63]** |
| Pre-subgenual frontal cortex | 0.59 ( 0.24 ) | 0.52 (0.2) | 0.74 (0.25) | **0.22 [0.14, 0.31]** |
| Cingulate gyrus, anterior | 0.55 ( 0.27 ) | 0.41 (0.19) | 0.8 (0.2) | **0.4 [0.32, 0.47]** |
| Inferior frontal gyrus | 0.44 ( 0.37 ) | 0.25 (0.21) | 0.79 (0.34) | **0.54 [0.43, 0.65]** |
| Superior frontal gyrus | 0.4 ( 0.34 ) | 0.2 (0.14) | 0.78 (0.27) | **0.58 [0.5, 0.67]** |
| Middle frontal gyrus | 0.34 ( 0.35 ) | 0.18 (0.19) | 0.64 (0.38) | **0.46 [0.34, 0.58]** |
| Postcentral gyrus | 0.32 ( 0.4 ) | 0.11 (0.11) | 0.68 (0.46) | **0.57 [0.42, 0.71]** |
| Superior parietal gyrus | 0.31 ( 0.35 ) | 0.14 (0.16) | 0.6 (0.39) | **0.46 [0.34, 0.59]** |
| Cingulate gyrus, posterior | 0.31 ( 0.27 ) | 0.21 (0.19) | 0.47 (0.3) | **0.26 [0.17, 0.36]** |
| Precentral gyrus | 0.23 ( 0.27 ) | 0.08 (0.09) | 0.48 (0.29) | **0.4 [0.3, 0.49]** |

**Baseline model**

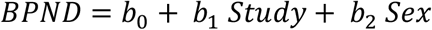

**Linear model**

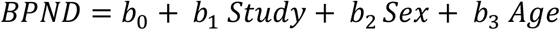

**Quadratic model**

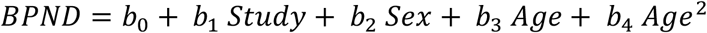

Model comparison was conducted contrasting these three regression models to each other within each region using the anova command in the R programming language, which tests the reduction in sum of squared error between models.

Percent difference per decade (PDD) was calculated using the following steps for each region: (1) a linear model with a single predictor (age) was fit to the data, (2) using the resulting regression equation, the estimated BP_ND_ at age 20 and age 30 were calculated, and (3) percent difference per decade was calculated using the following formula:

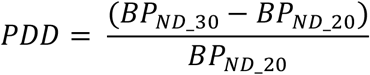

Confidence intervals for *PDD* were calculated by bootstrapping the above calculation using the boot package in R with 2000 repetitions (Canty & Ripley, 2017; Davison & Hinkley, 1997). A significance level of 0.05 can be inferred when zero is not contained within the 95% confidence interval.

Here we focus on complete reporting of effect sizes and confidence intervals rather than relying entirely on p-values which lead to somewhat arbitrary judgments of an effect being present or absent (Cumming, 2014). What has been the standard p-value heavy approach has been criticized by statisticians e.g. (Wasserstein & Lazar, 2016). The selection of a set of direct pairwise null hypothesis statistical tests would also be somewhat arbitrary given the large number of ROIs. For the comparisons across regions, we are focused on estimation and comparison of estimated effects (Gardner & Altman, 1986). Non-overlapping confidence intervals indicate a significant difference at p < .01 (Cumming, 2009). We highlight some differences between regions in the data, pointing out non-overlapping confidence intervals, but do not discuss every possible comparison. Note that direct comparisons between ROIs are not corrected for multiple comparisons.

## Results

### Average binding across regions of interest

Means and standard deviations of PVC-corrected BP_ND_ for each ROI are displayed in Table 2. As expected, BP_ND_ was highest in the striatum (ventral striatum: 37.16, putamen: 33.02, caudate: 26.9). The next highest BP_ND_ was observed in other medial and subcortical regions, but many of these values were an order of magnitude lower (pallidum: 14.83, subcallosal area: 9.23, insula: 2.44, thalamus: 2.43, amygdala: 3.02). The remaining frontal and temporal ROIs had mean BP_ND_ between 0.23 and 1.67.

### Relative strength of linear age effects

The largest raw age slopes (unstandardized coefficients from linear regression) were observed in striatal regions with smaller slopes in frontal and temporal regions with no age differences in subcallosal frontal cortex, pallidum, or hippocampus (Figure 1). However, since mean BP_ND_ differed by orders of magnitude across regions, unstandardized regression slopes (i.e., unit difference in BP_ND_ per year difference in age) were not directly comparable. The point estimates for each age slope (collapsing across but not controlling for sex and study) in each region were converted to percentage differences per decade and then qualitatively compared across regions (Figure 2). Estimated percentage differences in receptor BP_ND_ by age decade (linear effects) were highest in most temporal and frontal cortical regions (∼6–16% per decade), moderate in parahippocampal gyrus, pregenual frontal cortex, fusiform gyrus, caudate, putamen, thalamus, and amygdala (∼3–5%), and weakest in subcallosal frontal cortex, ventral striatum, pallidum, and hippocampus (∼0–2%). There were subcortical regions where age differences were relatively small and the upper bound of the 95% confidence intervals were less than 5% per decade: putamen, amygdala, thalamus, ventral striatum, hippocampus, and pallidum. In contrast, there were several cortical regions where age differences were relatively larger such that the lower bound of the 95% confidence intervals were greater than 5% per decade: postcentral gyrus, middle frontal gyrus, precentral gyrus, inferior, subgenual, and superior frontal gyri, anterior and posterior cingulate gyrus, superior parietal gyrus, inferiolateral parietal lobe, anterior and lateral orbital gyri, posterior and anterior superior temporal gyri, and lateral anterior temporal lobe. See non-overlapping confidence intervals in Figure 2.x

**Figure 1:**
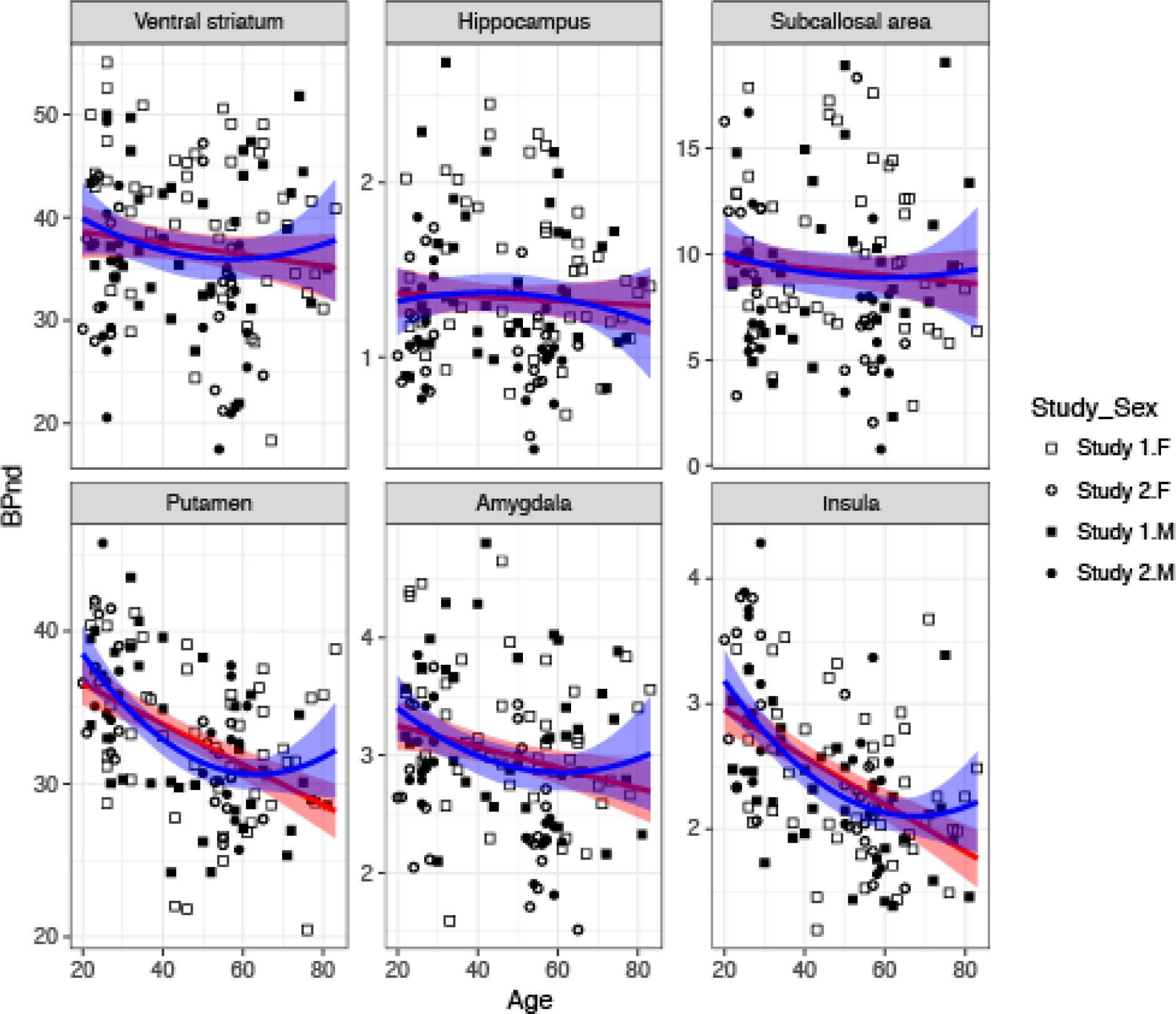
Linear and quadratic effects of Age on D2-BP_ND_ in select regions of interest. Pictures, scatterplots, and statistics for each ROI are available in an interactive app online at http://13.58.222.229:3838/agebp/.

**Figure 2:**
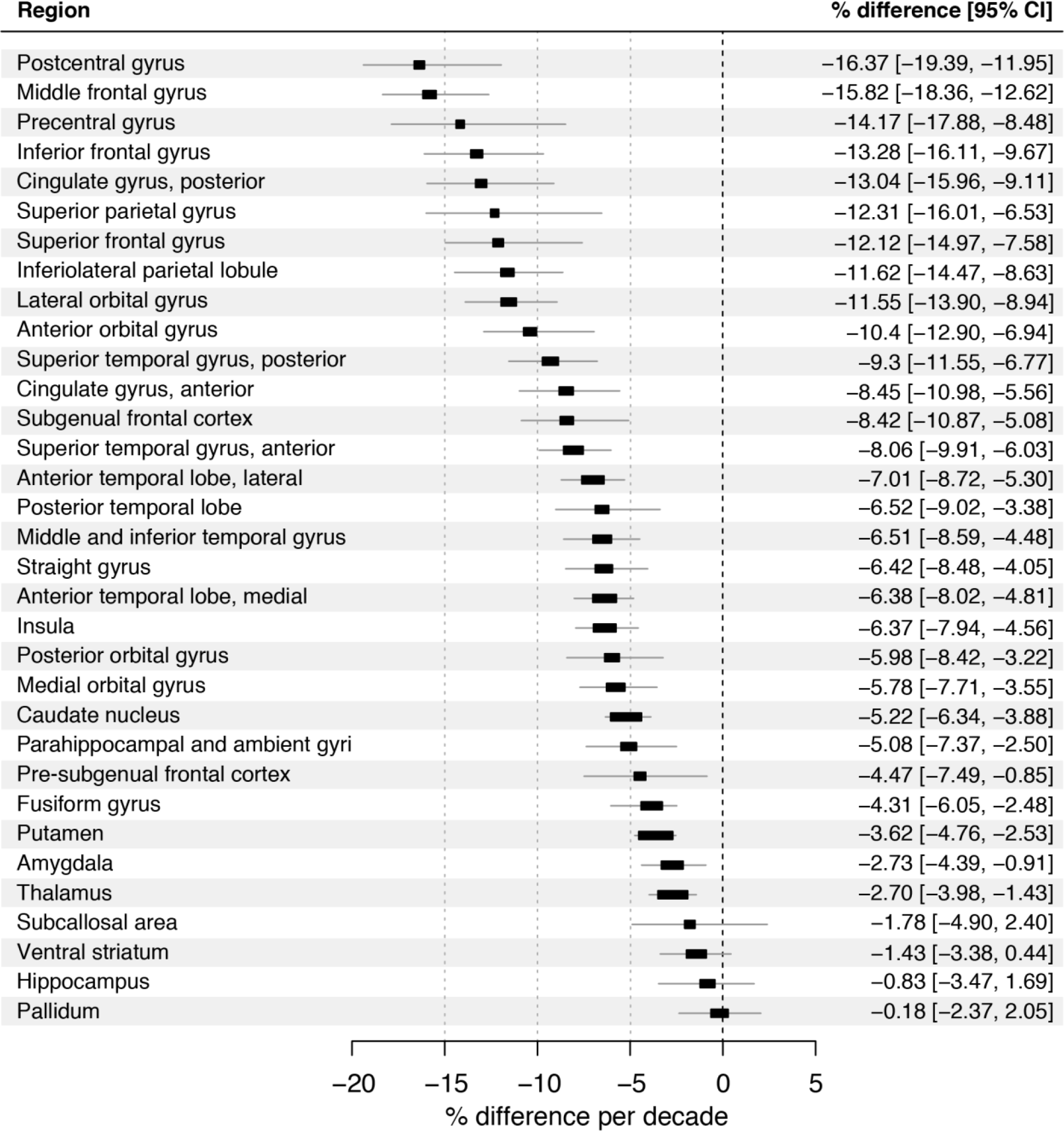
Percentage difference in D2-BP_ND_ per decade in regions of interest. Forest plot for all regions of interest. The position of the squares on the x-axis indicates the estimated percentage change in D2-BP_ND_ per decade and the horizontal bars indicate the 95% confidence intervals of the estimate. Non-overlapping confidence intervals indicate significant differences at p < .01 (Cumming, 2009).

### Non-linear effects of age

BP_ND_ in many frontal and temporal cortical regions were best fit by quadratic models (Tables 3–8). BP_ND_ in the straight gyrus/gyrus rectus, pre-subgenual frontal cortex, medial and posterior orbital gyri, fusiform gyrus, and all lateral temporal cortical regions showed concave-down quadratic effects of age such that BP_ND_ was reduced in middle age compared to young adulthood but then remained relatively stable (and low) until old age. Similar non-linear effects were observed in the insula and putamen such that there was a reduction in receptors during young adulthood (20–50 years old) that leveled off in middle age and older adulthood (+50 years old).

**Table 3.**
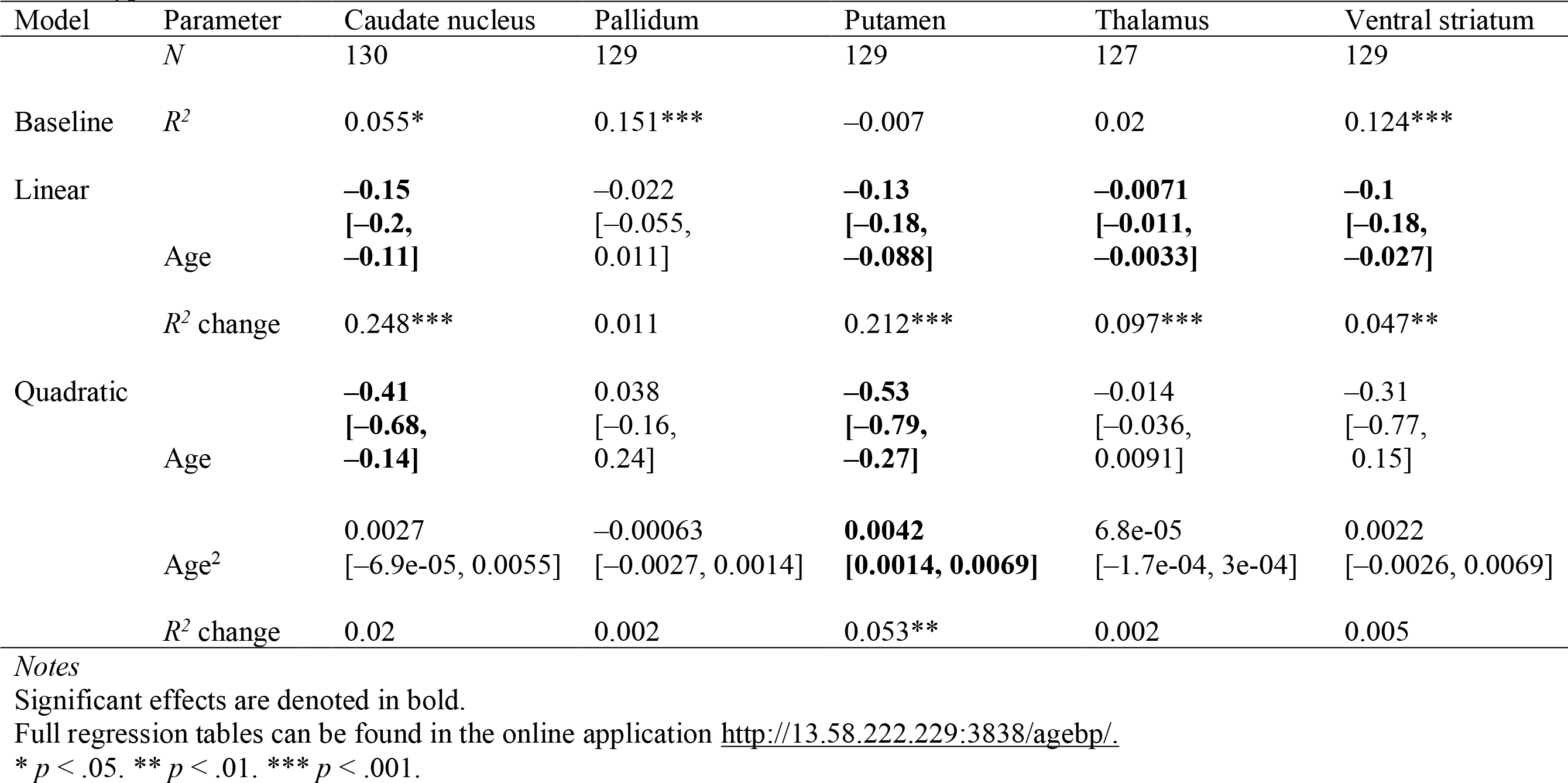
Multiple Linear Regression Analyses of Basal Ganglia ROI’s Age-related change in D2-like receptor availability (BP_ND_) using [^18^F]Fallypride PET.

| Model | Parameter | Caudate nucleus | Pallidum | Putamen | Thalamus | Ventral striatum |
| --- | --- | --- | --- | --- | --- | --- |
|  | *N* | 130 | 129 | 129 | 127 | 129 |
| Baseline | *R*<sup>2</sup> | 0.055* | 0.151*** | −0.007 | 0.02 | 0.124*** |
| Linear |  | **−0.15** | −0.022 | **−0.13** | **−0.0071** | **−0.1** |
|  | Age | [**−0.2,**<br>**−0.11**] | [−0.055,<br>0.011] | [**−0.18,**<br>**−0.088**] | [**−0.011,**<br>**−0.0033**] | [**−0.18,**<br>**−0.027**] |
|  | *R*<sup>2</sup> change | 0.248*** | 0.011 | 0.212*** | 0.097*** | 0.047** |
| Quadratic |  | **−0.41** | 0.038 | **−0.53** | −0.014 | −0.31 |
|  | Age | [**−0.68,**<br>**−0.14**] | [−0.16,<br>0.24] | [**−0.79,**<br>**−0.27**] | [−0.036,<br>0.0091] | [−0.77,<br>0.15] |
|  | Age<sup>2</sup> | 0.0027<br>[−6.9e-05, 0.0055] | −0.00063<br>[−0.0027, 0.0014] | **0.0042**<br>**[0.0014, 0.0069]** | 6.8e-05<br>[−1.7e-04, 3e-04] | 0.0022<br>[−0.0026, 0.0069] |
|  | *R*<sup>2</sup> change | 0.02 | 0.002 | 0.053** | 0.002 | 0.005 |

*Notes*
Significant effects are denoted in bold.
Full regression tables can be found in the online application http://13.58.222.229:3838/agebp/.
\* *p* < .05. \*\* *p* < .01. \*\*\* *p* < .001.

**Table 4.**
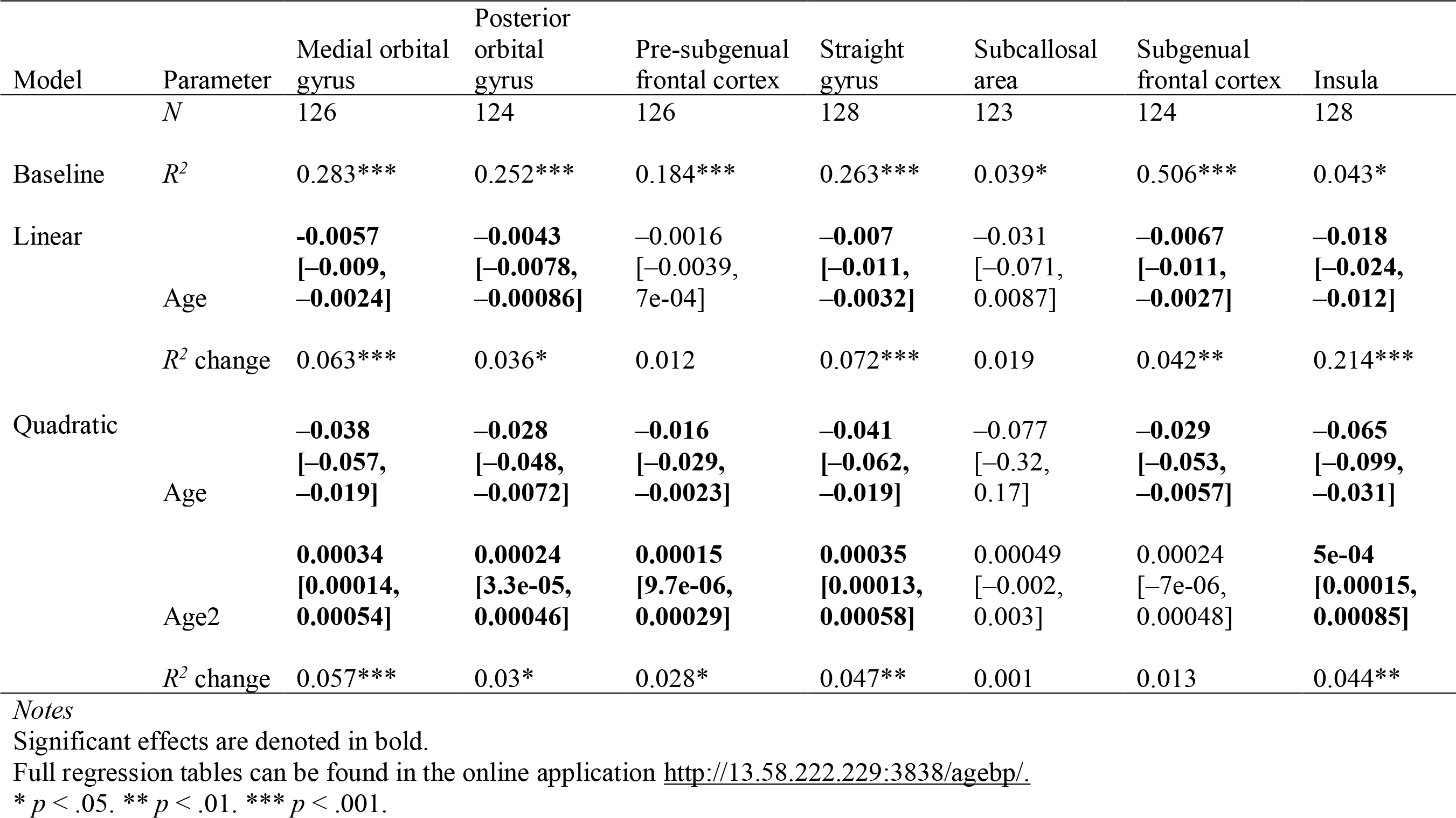
Multiple Linear Regression Analyses of Medial Frontal and Insula ROI’s Age-related change in D2–like receptor availability (BP_ND_) using [18F]Fallypride PET.

| Model | Parameter | Medial orbital gyrus | Posterior orbital gyrus | Pre-subgenual frontal cortex | Straight gyrus | Subcallosal area | Subgenual frontal cortex | Insula |
| --- | --- | --- | --- | --- | --- | --- | --- | --- |
|  | *N* | 126 | 124 | 126 | 128 | 123 | 124 | 128 |
| Baseline | *R*<sup>2</sup> | 0.283*** | 0.252*** | 0.184*** | 0.263*** | 0.039* | 0.506*** | 0.043* |
| Linear |  | **-0.0057** | **-0.0043** | -0.0016 | **-0.007** | -0.031 | **-0.0067** | **-0.018** |
|  | Age | **[-0.009, -0.0024]** | **[-0.0078, -0.00086]** | [-0.0039, 7e-04] | **[-0.011, -0.0032]** | [-0.071, 0.0087] | **[-0.011, -0.0027]** | **[-0.024, -0.012]** |
|  | *R*<sup>2</sup> change | 0.063*** | 0.036* | 0.012 | 0.072*** | 0.019 | 0.042** | 0.214*** |
| Quadratic |  | **-0.038** | **-0.028** | **-0.016** | **-0.041** | -0.077 | **-0.029** | **-0.065** |
|  | Age | **[-0.057, -0.019]** | **[-0.048, -0.0072]** | **[-0.029, -0.0023]** | **[-0.062, -0.019]** | [-0.32, 0.17] | **[-0.053, -0.0057]** | **[-0.099, -0.031]** |
|  | Age<sup>2</sup> | **0.00034**<br>**[0.00014, 0.00054]** | **0.00024**<br>**[3.3e-05, 0.00046]** | **0.00015**<br>**[9.7e-06, 0.00029]** | **0.00035**<br>**[0.00013, 0.00058]** | 0.00049<br>[-0.002, 0.003] | 0.00024<br>[-7e-06, 0.00048] | **5e-04**<br>**[0.00015, 0.00085]** |
|  | *R*<sup>2</sup> change | 0.057*** | 0.03* | 0.028* | 0.047** | 0.001 | 0.013 | 0.044** |

**Table 5.** Multiple Linear Regression Analyses of Anterior Frontal Lobe ROI’s Age-related change in D2-like receptor availability (BP_ND_) using [^18^F]Fallypride PET in Study 2.

| Model | Parameter | Anterior orbital gyrus | Cingulate gyrus, anterior | Inferior frontal gyrus | Lateral orbital gyrus | Middle frontal gyrus | Superior frontal gyrus |
| --- | --- | --- | --- | --- | --- | --- | --- |
|  | *N* | 129 | 130 | 128 | 125 | 124 | 121 |
| Baseline | R2 | 0.354*** | 0.48*** | 0.488*** | 0.284*** | 0.394*** | 0.683*** |
| Linear | Age | **-0.0063**<br>[ **-0.0093,**<br>**-0.0034** ] | **-0.0037**<br>[ **-0.0056,**<br>**-0.0018** ] | **-0.0059**<br>[ **-0.0085,**<br>**-0.0033** ] | **-0.0082**<br>[ **-0.011,**<br>**-0.005** ] | **-0.0068**<br>[ **-0.0095,**<br>**-0.0042** ] | **-0.0033**<br>[ **-0.0053,**<br>**-0.0013** ] |
|  | R2 change | 0.079*** | 0.052*** | 0.071*** | 0.122*** | 0.109*** | 0.027** |
| Quadratic | Age | **-0.035**<br>[ **-0.053,**<br>**-0.018** ] | **-0.014**<br>[ **-0.026,**<br>**-0.003** ] | **-0.035**<br>[ **-0.05,**<br>**-0.02** ] | **-0.038**<br>[ **-0.056,**<br>**-0.019** ] | **-0.034**<br>[ **-0.049,**<br>**-0.019** ] | **-0.023**<br>[ **-0.034,**<br>**-0.012** ] |
|  | Age2 | **0.00031**<br>[ **0.00012,**<br>**0.00049** ] | 0.00011<br>[ -5.9e-06,<br>0.00023 ] | **0.00031**<br>[ **0.00015,**<br>**0.00046** ] | **0.00031**<br>[ **0.00012,**<br>**5e-04** ] | **0.00028**<br>[ **0.00013,**<br>**0.00044** ] | **0.00021**<br>[ **8.9e-05,**<br>**0.00032** ] |
|  | R2 change | 0.045** | 0.013 | 0.049*** | 0.046** | 0.048*** | 0.027*** |

*Notes*
Significant effects are denoted in bold.
Full regression tables can be found in the online application http://13.58.222.229:3838/agebp/.
\* $p < .05$ . \*\* $p < .01$ . \*\*\* $p < .001$ .

**Table 6.** Multiple Linear Regression Analyses of Posterior Frontal and Parietal Lobe ROI’s Age-related change in D2-like receptor availability (BP_ND_) using [^18^F]Fallypride PET in Study 2.

| Model | Parameter | Cingulate gyrus, posterior | Precentral gyrus | Postcentral gyrus | Superior parietal gyrus | Inferiolateral remainder of parietal lobe |
| --- | --- | --- | --- | --- | --- | --- |
|  | *N* | 125 | 107 | 110 | 113 | 128 |
| Baseline | R2 | 0.224*** | 0.494*** | 0.465*** | 0.403*** | 0.31*** |
| Linear |  |  |  |  |  |  |
|  | Age | **-0.0046**<br>[ **-0.0069,**<br>**-0.0022** ] | **-0.0024**<br>[ **-0.0046,**<br>**-2e-04** ] | **-0.0062**<br>[ **-0.0093,**<br>**-0.0032** ] | **-0.0032**<br>[ **-0.006,**<br>**-0.00031** ] | **-0.0073**<br>[ **-0.011,**<br>**-0.0038** ] |
|  | R2 change | 0.081*** | 0.022* | 0.072*** | 0.025* | 0.083*** |
| Quadratic |  |  |  |  |  |  |
|  | Age | **-0.014**<br>[ **-0.029,**<br>**-0.00015** ] | -0.0096<br>[ -0.022, 0.0031 ] | **-0.035**<br>[ **-0.052,**<br>**-0.017** ] | **-0.021**<br>[ **-0.038,**<br>**-0.0043** ] | **-0.044**<br>[ **-0.064,**<br>**-0.025** ] |
|  | Age2 | 1e-04<br>[ -4.5e-05,<br>0.00025 ] | 7.6e-05<br>[ -5.6e-05,<br>0.00021 ] | **3e-04**<br>[ **0.00011,**<br>**0.00048** ] | **0.00019**<br>[ **1.4e-05,**<br>**0.00036** ] | **0.00039**<br>[ **0.00019,**<br>**0.00059** ] |
|  | R2 change | 0.011 | 0.006 | 0.04** | 0.023* | 0.062*** |

*Notes*
Significant effects are denoted in bold.
Full regression tables can be found in the online application http://13.58.222.229:3838/agebp/.
\* $p < .05$ . \*\* $p < .01$ . \*\*\* $p < .001$ .

**Table 7.** Multiple Linear Regression Analyses of Medial Temporal Lobe ROI’s Age-related change in D2-like receptor availability (BP_ND_) using [^18^F]Fallypride PET.

| Model | Parameter | Amygdala | Anterior temporal lobe medial part | Fusiform gyrus | Hippocampus | Parahippocampal and ambient gyri |
| --- | --- | --- | --- | --- | --- | --- |
|  | *N* | 131 | 130 | 127 | 129 | 127 |
| Baseline | R2 | 0.136*** | 0.169*** | 0.044* | 0.148*** | 0.198*** |
| Linear |  | **-0.013** | **-0.011** | **-0.0072** | -0.0037 |  |
|  | Age | **[-0.018, -0.0066]** | **[-0.015, -0.0069]** | **[-0.011, -0.0034]** | [-0.0078, 0.00038] | **-0.0033**<br>**[-0.0058, -0.00076]** |
|  | R2 change | 0.102*** | 0.162*** | 0.097*** | 0.021 | 0.041* |
| Quadratic |  | **-0.037** | **-0.032** | **-0.037** | 0.01 | **-0.018** |
|  | Age | **[-0.072, -0.0011]** | **[-0.054, -0.0091]** | **[-0.06, -0.015]** | [-0.014, 0.035] | **[-0.032, -0.0029]** |
|  | Age2 | 0.00026<br>[-0.00012, 0.00063] | 0.00022<br>[-1.4e-05, 0.00046] | 0.00032<br>[8.7e-05, 0.00055] | -0.00015<br>[-4e-04, 1e-04] | 0.00015<br>[-1.1e-06, 3e-04] |
|  | R2 change | 0.011 | 0.018 | 0.049** | 0.009 | 0.023 |

*Notes*
Significant effects are denoted in bold.
Full regression tables can be found in the online application http://13.58.222.229:3838/agebp/.
\* $p < .05$ . \*\* $p < .01$ . \*\*\* $p < .001$ .

**Table 8.** Multiple Linear Regression Analyses of Lateral Temporal Lobe ROI’s Age-related change in D2-like receptor availability (BP_ND_) using [^18^F]Fallypride PET.

| Model | Parameter | Superior temporal gyrus, anterior part | Anterior temporal lobe, lateral part | Middle and inferior temporal gyrus | Posterior temporal lobe | Superior temporal gyrus, posterior part |
| --- | --- | --- | --- | --- | --- | --- |
|  | *N* | 130 | 131 | 131 | 131 | 129 |
| Baseline | R2 | 0.194*** | 0.08** | 0.091*** | 0.111*** | 0.179*** |
| Linear | Age | **-0.012**<br>[-0.017, -0.0081] | **-0.01**<br>[-0.017, -0.0089] | **-0.0096**<br>[-0.014, -0.0055] | **-0.0051**<br>[-0.0084, -0.0019] | **-0.0086**<br>[-0.012, -0.0053] |
|  | R2 change | 0.163*** | 0.209*** | 0.128*** | 0.063** | 0.137*** |
| Quadratic | Age | **-0.048**<br>[-0.074, -0.023] | **-0.049**<br>[-0.074, -0.025] | **-0.055**<br>[-0.079, -0.032] | **-0.037**<br>[-0.056, -0.018] | **-0.043**<br>[-0.062, -0.023] |
|  | Age2 | **0.00038**<br>[0.00012, 0.00064] | **0.00038**<br>[0.00013, 0.00064] | **0.00048**<br>[0.00024, 0.00073] | **0.00034**<br>[0.00014, 0.00053] | **0.00036**<br>[0.00016, 0.00056] |
|  | R2 change | 0.039** | 0.046** | 0.082*** | 0.07*** | 0.06*** |

*Notes*
Significant effects are denoted in bold.
Full regression tables can be found in the online application http://13.58.222.229:3838/agebp/.
\* $p < .05$ . \*\* $p < .01$ . \*\*\* $p < .001$ .

## Discussion

This paper investigated regional differences in dopamine D2/3 (or D2-like) receptor BP_ND_ across the adult life span. The largest estimates of age-related differences per decade were observed in cortical regions (especially frontal and lateral temporal cortex), with more gradual loss of receptors in a subset of more medial cortical and subcortical regions. While we estimated declines of 6–16% in lateral temporal and many frontal cortical regions, we estimated striatal D2-like declines between 1.5 and 5% per decade. The estimates reported here are somewhat lower than those seen in a recent meta-analysis (Karrer et al., 2017), and could be due to our use of partial volume correction, an extremely healthy sample, or both. For instance, prior work from our lab on a subset of data from Study 1 noted that age-related changes in the uncorrected data more closely resemble those seen in the meta-analysis (Smith et al., 2017), while another study showed that compared to more sedentary adults, age-related change in striatal D2 BP_ND_ is less steep in physically active adults (Dang et al., 2017). Further, physical activity interventions have been shown to increase striatal D2 BP_ND_ (Robertson et al., 2016). Thus, the estimated changes reported here likely reflect both the partial volume techniques used and the relative health of our sample.

There was partial evidence for a medial/lateral distinction across cortical and subcortical regions. We found partial evidence for relative preservation of more ventromedial aspects of frontal cortex (subcallosal) and subcortical regions (ventral striatum, pallidum, hippocampus). This set of regions is partially consistent with components of the “core affect” functional network from studies of emotional processing (Lindquist, Wager, Kober, Bliss-Moreau, & Barrett, 2012) and the motivational loop described in anatomical and functional studies of dopaminergic frontostriatal circuits (Haber & Knutson, 2010; Seger & Miller, 2010). One exception is the moderate loss of receptors in the amygdala. One might expect more preservation in the amygdala relative to the hippocampus if the amygdala was primarily supporting affective function. Regardless, the relative preservation of dopaminergic function in the other medial regions may possibly account for the relative preservation of affective and motivational function with age.

There was inconsistent evidence for an anterior-posterior gradient. Although some anterior regions showed highest levels of percent difference per decade, several other posterior regions including parietal regions also showed large age differences relative to more anterior regions.

Although the steepest raw age slopes on BP_ND_ from the linear regressions were found in striatal regions, binding was also extremely high in these regions. Thus, even a large effect of age meant that the oldest adults still had relatively high levels of binding in striatum. This is why we chose to focus instead on the percentage age difference measures when making relative comparisons across regions. Although the percent difference per decade estimates did not control for study or sex (since these were dummy-coded categorical variables in the regression analyses), the slopes used to compute the percentage scores were nearly identical to the slopes from analyses that included the study and sex covariates. However, in contrast to the still relatively high levels of binding in older age in striatal regions, signal was low in most cortical regions in the oldest adults. This was the case especially for regions showing curvilinear age effects. In fact, the evidence for these curvilinear effects may be confounded somewhat by a floor effect given that measured BP_ND_ has a lower bound. An exception to this is the curvilinear effect is the putamen where the lowest values do not come close to approaching our threshold values. Portions of the putamen are connected to the pre/motor cortex and lateral prefrontal cortex. These corticostriatal loops are thought to mediate both motor and fluid cognitive abilities (Seger & Miller, 2010). Thus the curvilinear age differences in BP_ND_ in the putamen are consistent with both age-related motor slowing (Deary & Der, 2005) and age-related change in executive function and cognitive control (Rubin, 1999). However, smaller ROIs that isolated subareas more connected with lateral and motor cortex would be needed to fully evaluate this explanation.

Additionally, across the sample within our striatal ROIs we report higher BP_ND_ values in the ventral striatum compared to the putamen, whereas the opposite pattern has been reported using fallypride in previous studies using uncorrected data (i.e., no PVC) (Zald et al., 2010). Putamen BP_ND_ is higher than ventral striatum BP_ND_ in our uncorrected data as well (Study 1: ventral striatum BP_ND_ = 18.8, putamen BP_ND_ = 22.4). However, this relationship switches when using PVC. It is possible this occurs because the ventral striatum lies between ventricles and white matter; thus prior estimates of BP_ND_ in the ventral striatum likely included partial signal from neighboring ventricle and/or white matter in the estimates. Further, post-mortem studies comparing D2 receptor density in striatal subregions show a good deal of heterogeneity (Mawlawi et al., 2001), so it is not unreasonable for ventral striatal BP_ND_ to exceed putamen BP_ND_.

We found little evidence to support change in BP_ND_ in the hippocampus. This was somewhat unexpected, as studies of gray matter volume have documented both declines (Raz et al., 2010) and accelerated declines (Fjell et al., 2013) in both cross-sectional and longitudinal data. There is also some evidence for age-related decline in hippocampal dopamine receptors in human and non-human animals (Stemmelin et al 2000; Kaasinen et al 2000). However, the previous evidence from human PET imaging did not use partial volume correction; thus, age effects on BP_ND_ may have been somewhat confounded by age differences in gray matter volume. It is possible that there are age-related declines in hippocampal BP_ND_, but we were unable to detect them due to our sample size, restricted range in BP_ND_, and/or under-sampling of adults over the age of 65. In particular, given the greater variability in old-old age, future studies would benefit from over-sampling at the upper end of the human age range (Samanez-Larkin & D’Esposito, 2008). However, if there is true preservation of D2-like receptors in the hippocampus this would be an intriguing effect. Because memory performance has been linked to D2-like receptor binding in the hippocampus within older (Nyberg et al., 2016) and younger age (Takahashi et al., 2008) groups, age-related memory deficits have been suggested to be due to decline of the medial temporal lobes. However, these D2-like receptor effects have not been tested with cross-sectional age group or life-span designs. Our results may be viewed instead as consistent with the suggestion that many age-related memory deficits in healthy, disease-free adults are mediated by age-related changes in more frontal and/or lateral regions (Buckner, 2004).

There are important statistical caveats to the results reported here. In addition to study differences in average BP_ND_, there was a significant difference in the average age between the two studies (Study 1: *M* = 49.43 years old; Study 2: *M* = 41.40 years old). Thus, because study and age are correlated with each other and we controlled for study in our models, the slopes reported in this manuscript may under-estimate age effects. Also, the inflection points of many of the best-fitting quadratic models (e.g. in the majority of the frontal lobes, lateral temporal lobes, the fusiform gyrus and the insula) was in early middle-age (between ages 35-45). This is earlier than we predicted based on longitudinal studies of gray matter volume, which suggested the inflection point is in the mid-fifties (Raz et al., 2005), and corresponds to an age range not included in Study 2. Future studies are need to determine if these inflection points are true age effects, an artifact of floor effects (as mentioned above), and/or the result of under-sampling of this age range in these studies.

One major limitation of this study is that it was cross-sectional in nature. The estimates of age differences reported here (e.g., percentage difference per decade), and in the PET literature generally, are based on the assumption that cross-sectional studies accurately represent developmental trends. Until verified with longitudinal data, it remains possible that the age-related changes in dopaminergic function are at least partially a result of cohort effects.

Longitudinal studies of gray matter volume have estimated that cross-sectional studies can under-estimate the influence of aging (Raz et al., 2005) and similar longitudinal studies are necessary to determine to what extent this occurs in studies of the dopaminergic system. A longitudinal study is currently underway that should be able to address this question (Nevalainen et al., 2015). A second limitation of this study is that it focuses exclusively on D2-like receptor BP_ND_, without considering age-related changes in D1-like receptor binding, dopamine transporters, dopamine synthesis capacity, or dopamine release. Each of these measures a distinct aspect of dopaminergic function, and by focusing on D2-like receptors, we are likely to be missing important aspects of age-related change or stability elsewhere in the system. For example, D1-like receptor binding has shown steeper declines with age while synthesis capacity appears to remain stable with age (Karrer et al., 2017). However, it is important to note that the regional variation of these effects have not yet been systematically evaluated.

Collectively, the data presented here suggest that dopamine BP_ND_ does not show the same regional pattern of age-related changes observed in studies of gray matter volume and white matter integrity. There was somewhat surprising evidence for preservation of dopaminergic function well into older age in a subset of ventromedial cortical and subcortical brain regions. These results may help clarify one paradox of aging: namely, that some dopamine-mediated functions are preserved with age while others show marked decline. While there are clearly age-related differences in dopaminergic function across the adult life span, these changes in function are not uniform and they do not show the same regional pattern of change than has been observed using other neuroimaging methods. New theories of adult brain development are needed that incorporate these and other challenging results. Hopefully these findings inspire future studies that could be used to modify existing or propose new theories of human brain aging that better account for the differential changes in cognitive, affective, and motivational functions across adulthood.

## Acknowledgements

This study was supported by grants from the National Institute on Aging (R00-AG042596; T32-AG000029; R01-AG044838; R01-AG043458). We thank John Pearson, Duke University, for statistical consultation and advice as well as Kevin Seaman for web hosting consultation.

